# Hemidesmosomes regulate epidermal differentiation during embryogenesis

**DOI:** 10.1101/2025.10.01.679770

**Authors:** Juliet S. King, Kendall J. Lough, Scott E. Williams

## Abstract

In skin epidermis, integrins mediate adhesion of basal keratinocytes to the underlying basement membrane. While high expression of integrins has been correlated with stemness, there is limited direct evidence that integrins mediate keratinocyte retention within the basal layer. Here, we generate mosaic, epidermal-specific loss of integrin-β4 (encoded by *Itgb4*) or its ligand, laminin-α3β3ɣ2 (*Lama3*), using an *in utero* lentiviral-mediated approach. Although mutations in these genes cause postnatal skin blistering in mice and humans, we observe no evidence of epidermal-dermal separation embryonically. Despite no weobvious alterations to apicobasal polarity, *Itgb4*-deficient basal cells show mild defects in oriented cell divisions, with increased oblique divisions and altered telophase correction. However, differentiation via cellular delamination—where basal keratinocytes lose adhesion to the underlying basement membrane and transit into the suprabasal layer—is elevated upon *Itgb4* and *Lama3* loss. Notably, hyperactive Notch signaling both decreases integrin-β4 expression and increases delamination. These findings conclusively demonstrate a causal role for hemidesmosomes in regulating epidermal differentiation through both mitotic and non-mitotic mechanisms and shed additional light on the programs regulating delamination.

**Highlights:**

- Embryonic loss of integrin-β4 impairs hemidesmosome maturation, but does not cause blistering in the embryonic epidermis
- Integrin-β4 is dispensable for keratinocyte polarity but promotes telophase correction
- Hemidesmosome adhesions regulate differentiation through delamination

**Summary Statement:** Hemidesmosomes are specialized integrin complexes that anchor the epidermis to the basement membrane. We show that hemidesmosomal integrin-α6β4 and laminin-α3β3ɣ2 regulate epidermal differentiation by promoting basal cell retention in embryogenesis.

## INTRODUCTION

The epidermis is a stratified epithelium consisting of a mitotically active basal layer overlaid with suprabasal layers which differentiate via either asymmetric cell divisions (ACDs) or delamination. In an ACD, the spindle aligns along the apicobasal axis, displacing the apical daughter suprabasally, while during delamination cells exit the basal layer without dividing. Both ACDs and delamination contribute to epidermal differentiation during embryogenesis, with delamination driving initial stratification and ACDs peaking later (Damen et al., 2021; Descovich et al., 2023; Lough et al., 2020; Williams et al., 2014). In adulthood, ACDs are uncommon and differentiation is driven mainly by delamination (Cockburn et al., 2022; Ipponjima et al., 2016; Liu et al., 2019; Rompolas et al., 2016). Despite evidence of delamination through *ex vivo* live imaging and clonal lineage tracing, little is known about what regulates this process *in vivo* (Cockburn et al., 2022; Damen et al., 2021; Ellis et al., 2019; Mesa et al., 2018; Miroshnikova et al., 2018).

Basal keratinocytes adhere to an underlying extracellular matrix (ECM)-rich basement membrane via two major classes of integrins. Integrin-α6β4 dimers complex with the cytokeratin intermediate filament network to form hemidesmosomes (HDs), while α3β1 dimers integrate into focal adhesions and link to the actin cytoskeleton (Rousselle et al., 2022). The importance of HDs is exemplified by the severe blistering disease that occurs in mice and humans upon disruption of HD components or their ligand, laminin-α3β3ɣ2 (laminin-332), known as junctional epidermolysis bullosa (JEB) (Dowling et al., 1996; Pulkkinen and Uitto, 1998; Raymond et al., 2005). While HDs are viewed as stronger and more stable than focal adhesions, emerging evidence suggests HDs can modulate dynamic processes such as migration, force generation and keratin fiber assembly (Löffek et al., 2014; Moch and Leube, 2021; Wang et al., 2020).

High levels of integrin-β1 have been linked to keratinocyte stemness, while loss of surface integrin-β1 and integrin-α6 are correlated with differentiation (Adams and Watt, 1990; Jones et al., 1995; Li et al., 1998). Paradoxically, expression of differentiation markers precedes loss of surface integrins (Adams and Watt, 1990; Cockburn et al., 2022), raising the question of whether integrin loss is cause or consequence of differentiation. Adding to this complexity, mice lacking components of either integrin complex show no obvious defects in epidermal differentiation, even in non-blistered and embryonic skin (DiPersio et al., 2000; Dowling et al., 1996; Georges-Labouesse et al., 1996; van der Neut et al., 1996).

Here, we analyze the consequences of mosaic *Itgb4* or *Lama3* loss in the developing epidermis and show that—although epidermal architecture is surprisingly normal with no signs of overt blistering—mutant cells are more prone to differentiate than their wild-type (WT) neighbors. Mitotic *Itgb4* knockdown keratinocytes have subtle spindle orientation defects, with a trend toward increased persistent oblique divisions. Most notably, delamination is markedly increased upon loss of *Itgb4* or its ligand *Lama3*. Finally, we show that Notch signaling, an epidermal differentiation regulator, inhibits integrin-β4 expression and promotes delamination. Collectively, these studies demonstrate a role for HDs in promoting basal layer retention and provide mechanistic insights into the pathways that regulate delamination.

## RESULTS AND DISCUSSION

Since germline or whole epidermal loss of *Itgb4* results in dermal-epidermal separation (Dowling et al., 1996; Raymond et al., 2005), potentially confounding interpretation of architectural defects, we utilized an *in utero* lentiviral-mediated approach to generate mosaic *Itgb4* knockdown (Fig. 1A). Using the LUGGIGE (lentiviral ultrasound-guided gene expression and gene inactivation) technique (Beronja et al., 2010), transduction of progenitor basal keratinocytes occurs at embryonic day (E)9.5, prior to stratification, with RFP-positivity as a reporter of infection. RFP+ basal cell phenotypes can then be compared to both uninfected littermates (WT) and neighboring, uninfected cells (RFP-). *Itgb4^4124^* and *Itgb4^2326^* RFP+ cells showed robust depletion of *Itgb4* at the mRNA level *in vitro*, and of integrin-β4 antibody intensity at the dermal epidermal junction (DEJ) *in vivo* (Fig. 1B-D; Fig. S1A-C). Of note, reduction of integrin-β4 was cell-autonomous and localized to regions of RFP+ basal cells.

**Figure 1.**
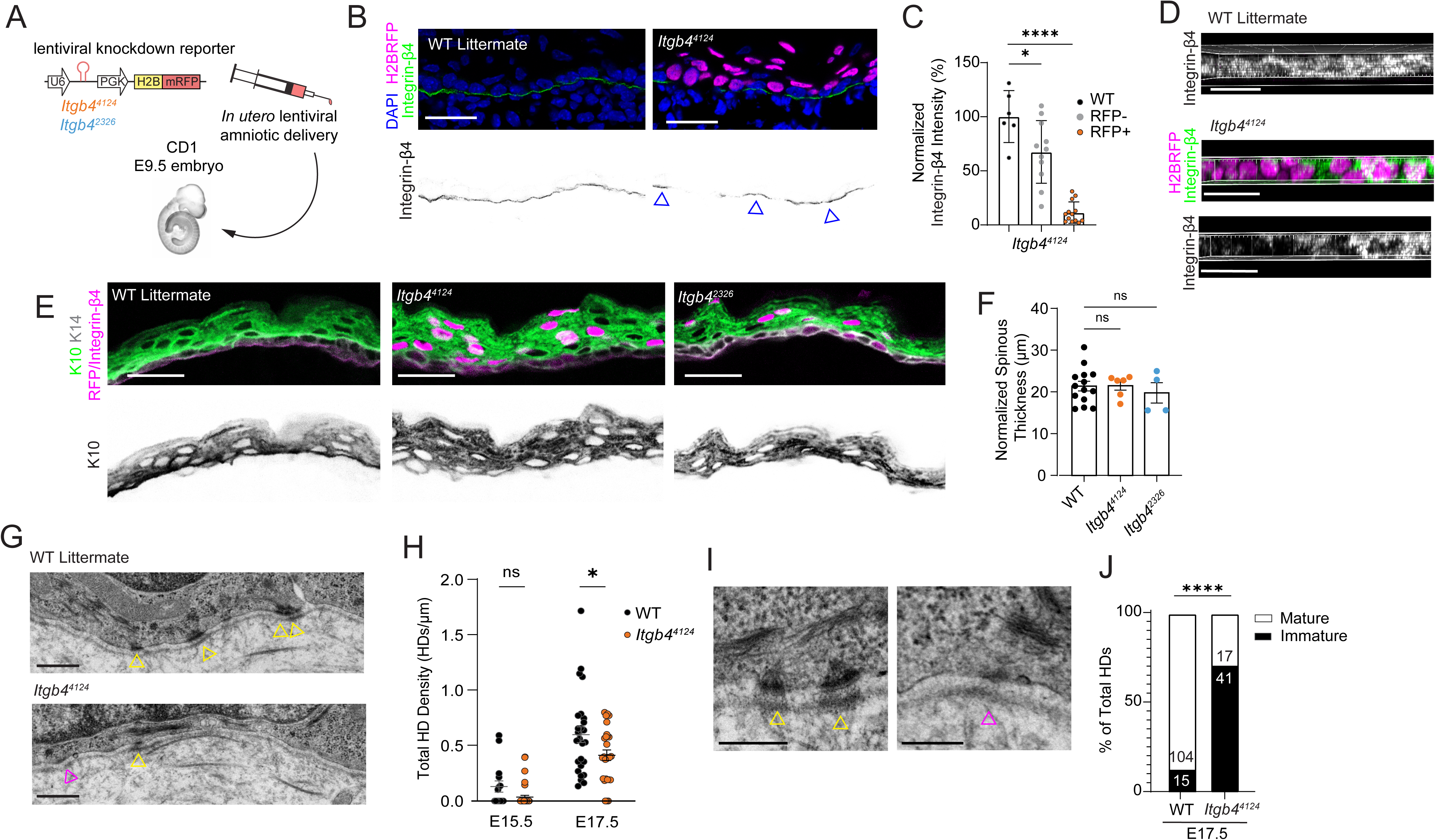
Mosaic loss of *Itgb4* alters embryonic basement membrane organization without overt blistering. (A) Schematic of LUGGIGE technique. (B-D) *In vivo* validation of *Itgb4^4124^*shRNA. Single-plane confocal images (B) and *en face* z-stack reconstruction (D) of E17.5 mosaic epidermis showing RFP+ knockdown cells and integrin-β4, with quantification of integrin-β4 fluorescent intensity (C). Blue arrowheads in (B) represent areas of non-transduced (RFP-) basal cells. (E,F) Single-plane confocal images (E) and quantification (F) of cytokeratin-10 (K10) immunofluorescence in E16.5-17.5 control, *Itgb4^4124^* and *Itgb4^2326^* epidermis; each dot in (F) represents average thickness per animal. (G-J) Transmission electron micrograph images (G,I) and quantification (H,J) of hemidesmosomes (HD) in control and highly-transduced *Itgb4^4124^* nape skin. Total HD density is quantified in (H) and maturity in (J); yellow arrowheads indicate mature and magenta arrowheads immature HDs. Scale bars: 25µm (B, D, E), 0.5µm (G), and 0.25µm (I). p-values: ns, not significant; * p<0.05, **** p<0.0001.

*Itgb4^4124^* and *Itgb4^2326^* epidermis had equivalent thickness of the suprabasal (K10+) layers compared to controls (Fig. 1E,F). Although overt signs of blistering in these animals were lacking, we utilized transmission electron microscopy to examine the DEJ at an ultrastructural level. HDs were sparse at E15.5 in both the *Itgb4^4124^*and age-matched WT DEJ (Fig. 1G). However, in agreement with previous studies in neonatal mice, E17.5 *Itgb4^4124^* epidermis showed reduced HD density compared to WT controls (Dowling et al., 1996; Raymond et al., 2005; van der Neut et al., 1996) (Fig. 1H). While transduction rates are highest in nape skin, residual HDs in *Itgb4^4124^* sections could be explained by infection mosaicism. We also observed variability in the morphology of the HDs, which included 1) mature HDs, which are roughly triangular, electron dense, contain anchoring fibrils extending into the lamina densa and clear keratin filament attachments intracellularly, and 2) immature HDs, which lack one or more of these defining characteristics (Guo et al., 1995). In addition to a decrease in the overall number of HDs observed at the DEJ, *Itgb4^4124^* tissue had a significantly greater proportion of immature HDs (Fig. 1J; S1D,E). These data indicate that loss of integrin-β4 contributes to DEJ instability independent of tissue stress.

Although *Itgb4* knockdown epidermis showed normal expression of differentiation markers, we noticed that RFP+ cells appeared more abundant in suprabasal layers, suggesting an increased propensity to differentiate (Fig. 2A). Taking advantage of the inherent mosaicism of LUGGIGE, we reasoned that all RFP+ spinous cells are the result of either an asymmetric division or delamination event. In this way, RFP-positivity serves as a surrogate lineage trace, and is particularly effective when transduction levels are at low “clonal” density. We defined the “differentiation index” as the proportion of suprabasal cells which are RFP+ divided by the proportion of basal cells which are RFP+. Compared to *Scramble* controls, which had a differentiation index near one (1.032), *Itgb4^2326^*and *Itgb4^4124^* epidermis ratios skewed higher (1.521 and 1.212, respectively; Fig. 2B-D). These data show that loss of integrin-β4 leads to a comparative disadvantage in competing for basal layer occupancy, favoring differentiation.

**Figure 2.**
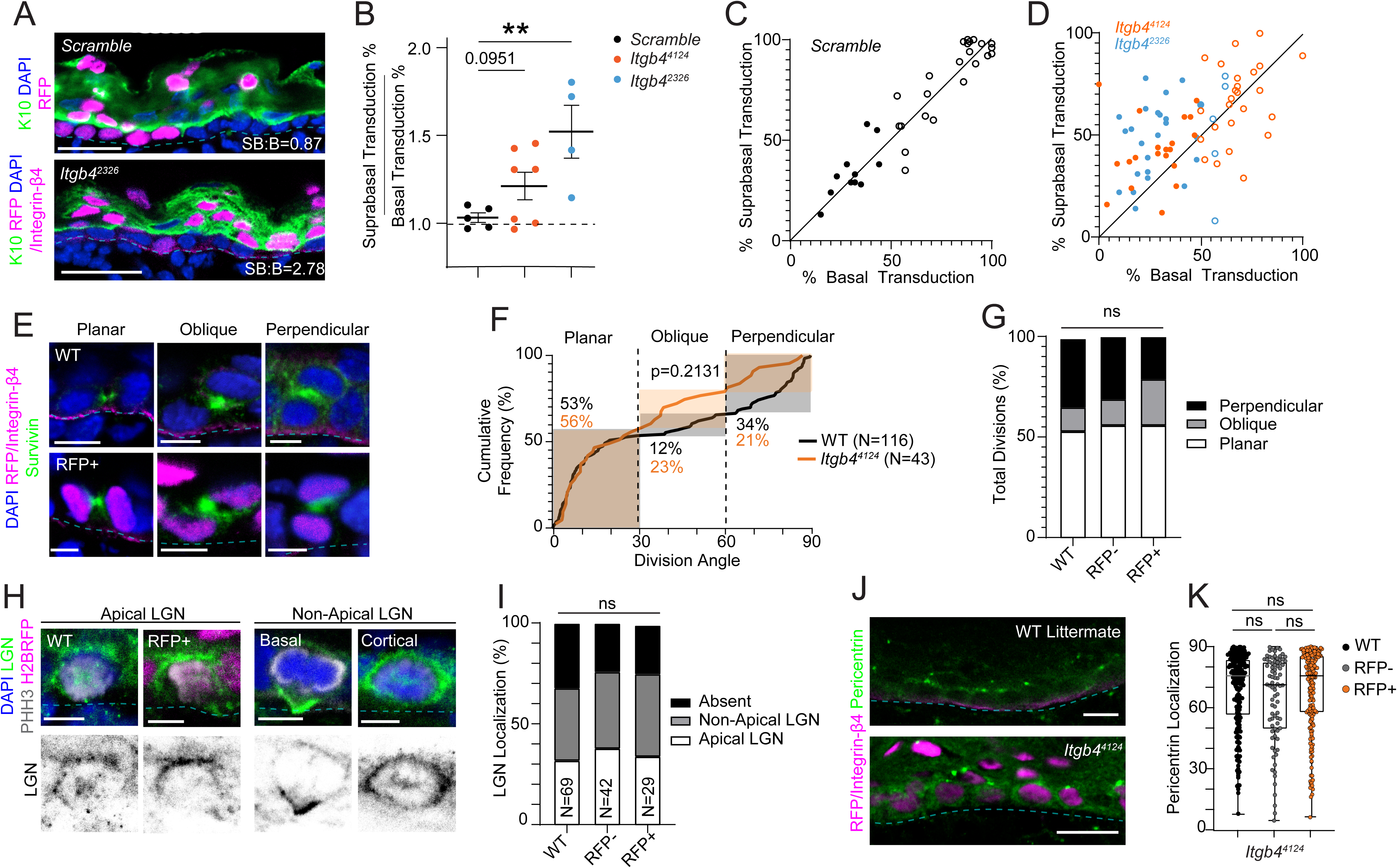
*Itgb4* loss leads to decreased basal occupancy but does not impact division orientation or polarity. (A) Single-plane confocal images of *Scramble* and *Itgb4^2326^*epidermis. (B-D) Differentiation index values for *Scramble*, *Itgb4^4124^* and *Itgb4^2326^* epidermis. Average ratios for each animal plotted in (B), with specific basal and suprabasal transduction values for individual images shown in (C,D); values greater than 50% basal infection (saturation density) represented with open circles. (E) WT and *Itgb4^4124^*RFP+ telophase cells (survivin, green) in planar, oblique, and perpendicular orientations. (F) Cumulative frequency plot of WT (black) and *Itgb4^4124^* (orange) division orientation angles; Kolmogorov-Smirnov cumulative frequency test, p=0.2131. (G) Bar graph of orientation types, binned as by shaded bars in (F). (H,I) Examples of LGN staining patterns, quantified in (I). (J,K) Images of pericentrin localization in WT and *Itgb4^4124^* epidermis (J), quantified in (K), where apical is 90°. Scale bars: 10μm in (E, H); 25μm in (A, J). DEJ indicated by dashed cyan lines. p-values: ns, not significant; determined by t-test, except (G, I), chi-square, and (K), one-way ANOVA.

A reduction in RFP+ cells relative to RFP-cells in the basal layer could be explained by hypoproliferation, but we did not observe differences in expression of either the mitotic marker phospho-histone H3 (pHH3) or incorporation of 5-Ethynyl-2-deoxyuridine (EdU) over a two-hour window (Fig. S2A-D). Additionally, there was no evidence of elevated cell death upon *Itgb4* knockdown, where apoptotic (cleaved-caspase 3+) cells were rare, and not more frequent in RFP+ cells (Fig. S2E,F). Next, we asked whether a shift in division orientation could explain the basal occupancy deficit. During peak stratification (E16.5-E17.5) there are roughly equal proportions of planar symmetric cell divisions and perpendicular differentiative ACDs, with very few oblique-oriented divisions (Lough et al., 2019). Using the mid-body marker survivin, we measured the division angle of telophase cells relative to the basement membrane in RFP+, RFP- and WT littermate basal cells (Fig. 2E). In both *Itgb4^4124^* and *Itgb4^2326^* epidermis, we observed an atypical, but mild increase in oblique divisions at the expense of perpendicular divisions (Fig. 2F,G; S2G-H). However, since the proportion of planar divisions was similar in control (RFP- or WT) and *Itgb4* knockdown (RFP+) cells, it seems unlikely that decreased self-renewal could explain the relative paucity of basal cells.

Division orientation is regulated in two steps in the developing epidermis. First, during pro-metaphase, spindle orientation complex proteins—including Par3 and LGN—promote perpendicular orientation, while antagonists like AGS3 inhibit LGN function to maintain planar divisions (Descovich et al., 2023; Lechler and Fuchs, 2005; Lough et al., 2019; Williams et al., 2014). Later, persistent obliques resolve to planar or perpendicular in a process called telophase correction (Descovich et al., 2023; Lough et al., 2019). As in germline-deleted *Itgb4* epidermis (Lechler and Fuchs, 2005), we observed no change in *Itgb4^4124^*RFP+ basal cells with polarized apical LGN crescents (Fig. 2H,I). Initial spindle positioning requires intact epithelial polarity (Vorhagen and Niessen, 2014; Williams et al., 2014), but we found no alterations to the apical localization of the polarity protein Par3 (Fig. S2I-K) or the apical positioning of centrosomes (Fig. 2J,K). These data suggest that the mild increase in oblique divisions cannot be explained defects in spindle orientation.

To assess telophase correction behavior, we performed *ex vivo* live imaging on mosaically-transduced *Itgb4^4124^*E16.5 epidermal explants (Cetera et al., 2018; Descovich et al., 2023; Lough et al., 2019). We used mice on a *Krt14^Cre^;Rosa26^mTmG^* background such that epidermal keratinocytes are labeled with membrane-GFP while the underlying epidermis is labeled with membrane-Tomato (Fig. 3A). Explants were imaged *en face*, and the division angle was measured relative to the underlying DEJ via z-reconstructions from anaphase onset (time, t=0) to 60 minutes post-anaphase (t=60; hereafter referred to as telophase; Fig. 3B,C). Generally, both RFP- and RFP+ cells which entered anaphase at planar or perpendicular orientations remained in these positions through telophase (Fig. 3C-F). However, telophase obliques were more frequent among RFP+ cells (12%) compared to RFP-cells (3%). Moreover, the majority (55%) of RFP+ cells that enter anaphase at an oblique angle remained obliquely positioned, a behavior which was never observed in RFP-cells (Fig. 3E). Previously, we showed that anaphase oblique cells correct to planar when the apical daughter retains basement membrane contact via a “basal endfoot” (Lough et al., 2019). Interestingly, RFP+ obliques were observed which retained an endfoot for 60 minutes or more yet failed to planar correct (Fig. 3G). These observations suggest that the mild increase in oblique divisions observed in fixed tissue is likely attributable to impaired telophase correction.

**Figure 3.**
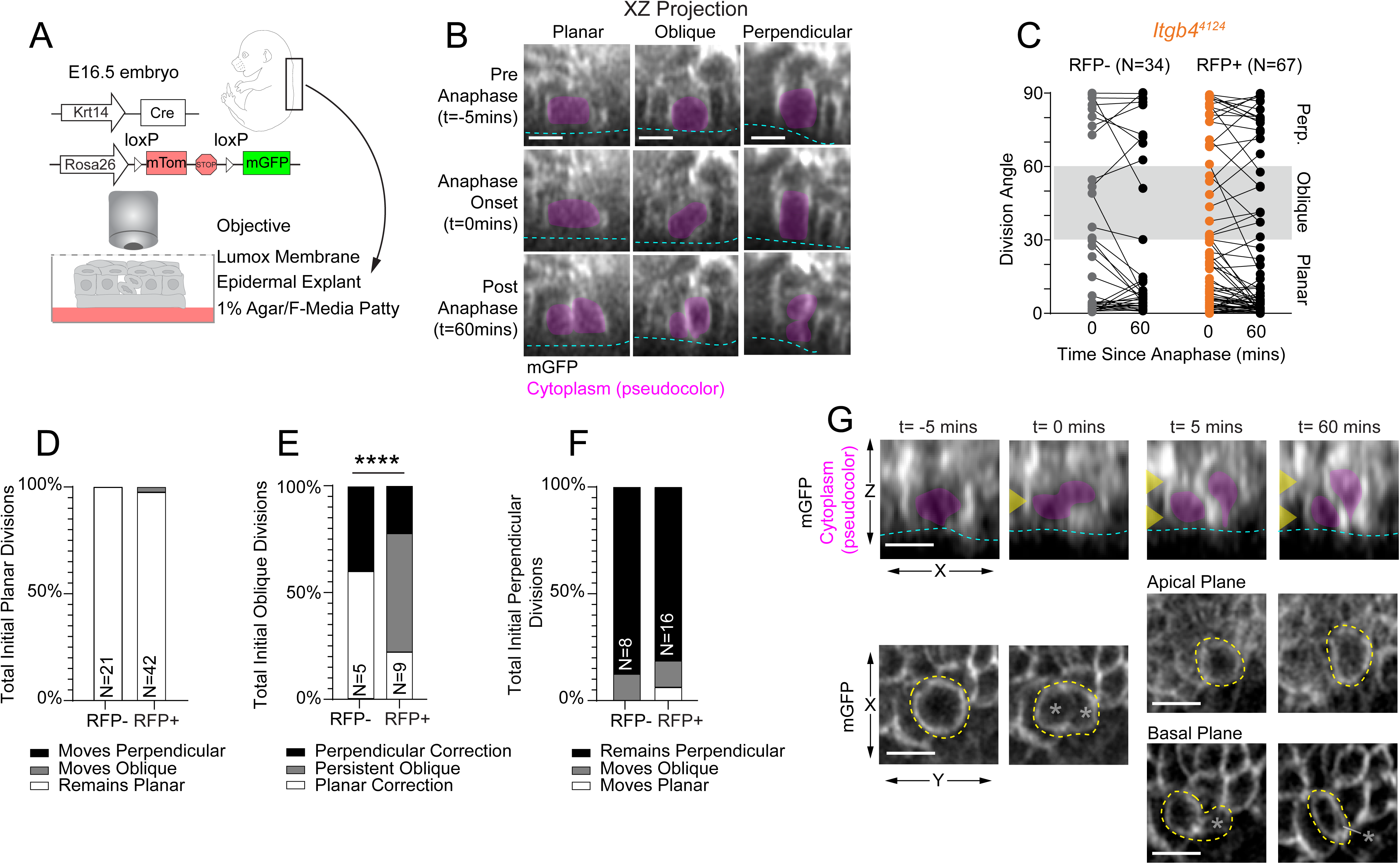
Failure of telophase correction in *Itgb4* KD cells contributes to increase in oblique divisions. (A) Schematic of *ex vivo* live imaging. (B) XZ projections of dividing cells from pre-anaphase (t=-5mins), anaphase (t=0mins), and post-anaphase timepoints (t=60mins). Long axis (psuedocolored magenta) of the basal cells membrane (mGFP, white) used to determine division angle relative to the underlying dermis. (C) Line graphs of division orientation angle per cell between anaphase and 60mins post-anaphase in *Itgb4^4124^* RFP-(grey) and RFP+ cells (orange). (D-F) Bar graph of data from (C) depicting the correction behavior of initial planar (D), oblique (E), and perpendicular (F) RFP- versus RFP+ cells. (G) Representative image of an initial oblique-oriented cell with persistent basal endfoot. Yellow arrows indicate the optical sections in XZ corresponding to the apical and basal planes below. p-values: **** p<0.0001 by observed vs. expected chi-square test in (D-F). Scale bars:10μm.

In addition to ACDs, basal keratinocytes can differentiate through a division-independent mechanism termed delamination. While poorly understood at a molecular level, delaminating cells initiate expression of differentiation markers while still in the basal layer (Cockburn et al., 2022; Damen et al., 2021; Williams et al., 2014). Thus, to measure delamination, we quantified the frequency of Keratin-14 (basal keratin) and Keratin-10 (spinous keratin) dual-positive basal cells which retained basement membrane contact (Fig. 4A) (Williams et al., 2014). At E16.5-E17.5, compared to RFP- and WT controls, both *Itgb4^4124^* and *Itgb4^2326^* RFP+ showed an increased propensity for dual-positive basal cells (Fig. 4B). Although RFP+ cells transduced with a non-targeting *Scramble* control shRNA were slightly more prone to delaminate than non-transduced RFP-cells (7.1% vs 4.8% of basal cells), *Itgb4^4124^* and *Itgb4^2326^* RFP+ cells showed significantly higher dual-positivity rates (13.4%, 14.8%). This phenotype was age-specific, because no increase in delamination frequency was observed in RFP+ *Itgb4^4124^*basal cells at E15.5 (Fig. S4A). This observation agrees with our finding that there are fewer, and less mature, HDs at E15.5 compared to E17.5 (Fig. 1H; S1D,E).

**Figure 4.**
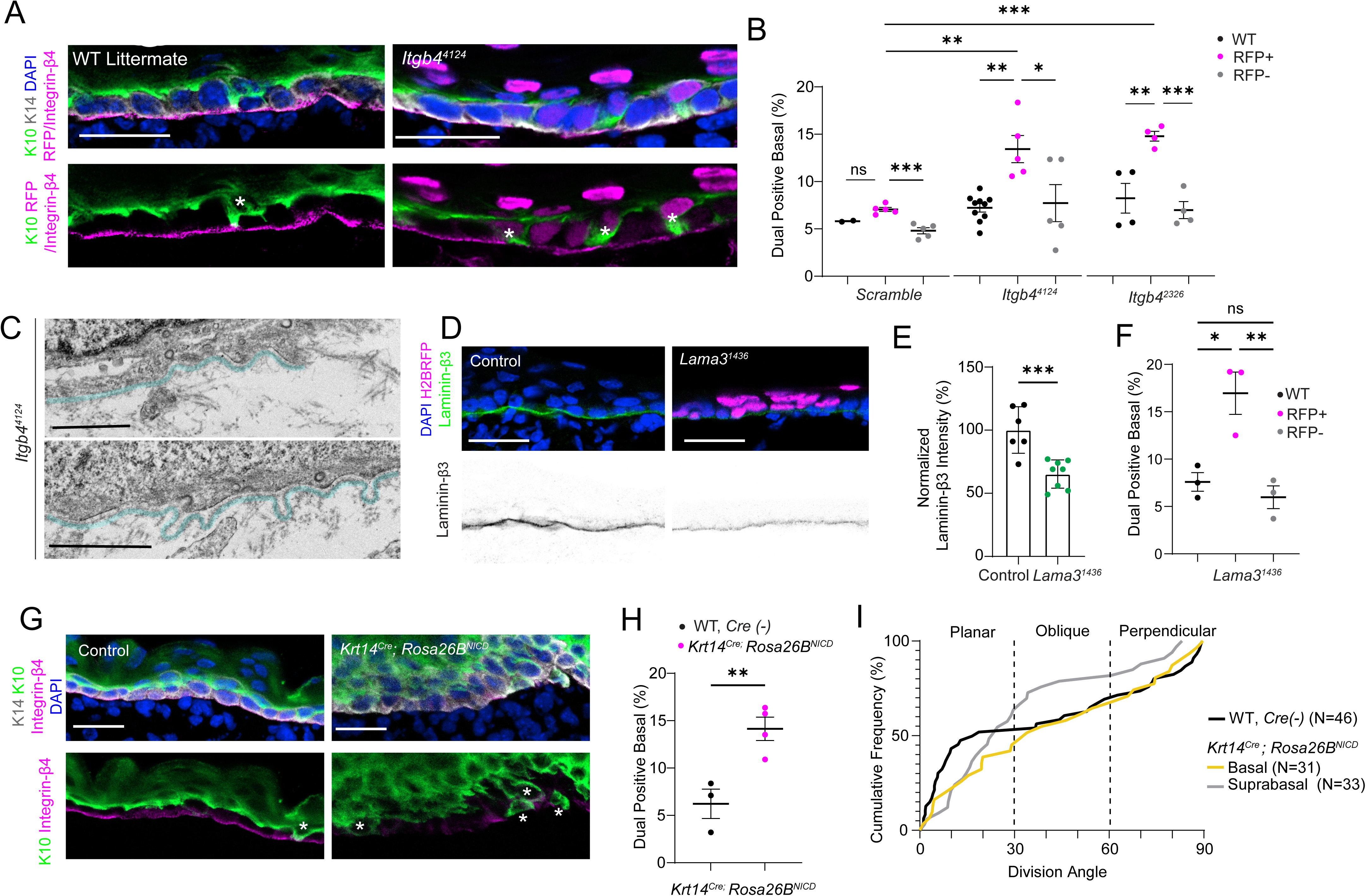
Loss of integrin-β4 or its ligand laminin-332 cell-autonomously increases delamination. (A) Control and *Itgb4^4124^* knockdown epidermis at E17.5, stained with spinous marker Keratin-10 and basal marker Keratin-14. (B) Quantification of dual-positive cells for WT littermates, RFP+ and RFP-populations in *Scramble*, *Itgb4^4124^* and *Itgb4^2326^* backskins; each dot represents one animal. (C) TEM micrograph of *Itgb4^4124^* mutant epidermis with aberrant laminae densa architecture (pseudocolored in cyan). (D,E) Representative images (D) of laminin-β3 intensity in control and *Lama3^1436^*epidermis, quantified in (E). (F) Dual-positive basal cell frequency for *Lama3^1436^*mutant epidermis, as in (B). (G,H) Images of E16.5 *Krt14^Cre^;Rosa26B^NICD^* and Cre-negative control epidermis (G) stained as in (A), quantified in (H). Asterisks in (A,G) indicate dual-positive basal cells, inferred to be delaminating. Scale bars: 25μm in (A, D, G); 1μm in (C). p-values by t-test: ns, not significant; * p<0.05; ** p<0.01; *** p<0.001.

Integrin-α6β4 dimers bind to laminin-332, which is produced and secreted by basal keratinocytes (Rousselle and Beck, 2013; Rousselle et al., 2022). In addition to decreased maturity and number of mature HDs, *Itgb4^4124^* tissue showed loops and occasional breaks in the lamina densa (pseudocolored cyan in Fig. 4C). Moreover, immunofluorescent staining for the laminin-β3 subunit of laminin-332 appeared weaker in *Itgb4* knockdown epidermis compared to littermate controls (Fig. S4B), suggesting that laminin-332 may also be an important regulator of differentiation behavior. To test this, we designed an shRNA, *Lama3^1436^*, to target one of its subunits. As validation of its efficacy *in vivo*, we observed a 35% reduction of Laminin-β3 intensity at the DEJ, indicating successful reduction of the laminin-332 trimer (Fig. 4D,E). Although *Lama3* loss did not affect epidermal thickness, wholemount imaging revealed a dramatic increase in RFP+ cells in the suprabasal layers relative to the basal layer (Fig. S4C,D). Indeed, the *Lama3^1436^*RFP+ differentiation index (2.040, p=0.0084) was even higher than observed upon *Itgb4* loss. Finally, K10/K14 dual-positive basal cells in *Lama3^1436^*epidermis were enriched in the RFP+ population compared to RFP- (17.0% vs. 6.0%; Fig. 4F; S4E). Taken together, these data show that disruption of either HD receptors or their ligand increases the rate of delamination *in vivo*. To shed additional light on the mechanisms regulating delamination, we turned to a well-known regulator of epidermal differentiation, Notch signaling (Nowell and Radtke, 2013; Watt et al., 2008). Developmentally, Notch is considered pro-differentiative because conditional deletion of its transcriptional effector *Rbpj* results in impaired differentiation and epidermal thinning, while overexpression of the active Notch1 intracellular domain (NICD1) increases spinous thickness (Blanpain et al., 2006; Williams et al., 2011). We previously showed that *Rbpj* loss does not affect asymmetric cell divisions (Williams et al., 2011), suggesting that Notch’s role in differentiation may be through delamination. To test this, we utilized *Krt14^Cre^;Rosa26B^NICD^*mice which overexpress active Notch1 (NICD1). Of note, these mice show reduced hemidesmosomes and loss of integrin-β4 at the DEJ (Blanpain et al., 2006), which we independently confirmed (Fig. 4G; S4F). Quantification of K10/K14+ dual-positive basal cells revealed a significant increase in delamination upon NICD1 overexpression, with little change in division orientation (Fig. 4G-I). Therefore, although NICD1 overexpression also increases suprabasal mitoses (Fig. 4I, gray line), elevated delamination is likely to be a primary contributor to the tissue-wide, hyper-differentiation seen in Notch overexpression mice.

Basal residency of keratinocytes is a key determinant of their progenitor capacity, with the expression level of integrins correlating with stemness (Jensen et al., 1999; Jones and Watt, 1993; Jones et al., 1995; Li et al., 1998). Despite this long-standing association, mice lacking one or multiple integrin subunits typically display the blistering that is a hallmark of junctional epidermolysis bullosa, but have otherwise normal epidermal architecture (Dowling et al., 1996; Raghavan et al., 2000; Raymond et al., 2005). This raises the question of if, and how, integrins regulate epidermal differentiation. Here, we provide evidence that loss of *Itgb4* or its ligand *Lama3* lead to increased delamination and abnormal telophase correction, demonstrating a key role for HDs in promoting basal layer retention.

Classically, delamination has been viewed as the major driver of epidermal differentiation (Watt, 1984; Watt and Green, 1982), but in recent years it has become clear that both ACDs and delamination play critical, but context-dependent, roles (Smart, 1970; Williams et al., 2011). While perpendicular divisions are a key driver of differentiation during peak stratification, they are not required for initial stratification or adult homeostasis in skin, although ACDs persist into adulthood in other stratified epithelia (Byrd et al., 2019; Ipponjima et al., 2016; Lechler and Fuchs, 2005; Rompolas et al., 2016; Williams et al., 2011). Here, we show that cell-matrix adhesions also play a minor role in regulating telophase correction, adding to the list of extrinsic factors such as tissue geometry, apoptosis, and cell-cell adhesions that influence division orientation (Box et al., 2019; Lough et al., 2019; Soffer et al., 2022).

Although direct evidence of embryonic delamination is limited, it likely occurs on a timescale of minutes to hours, while in the adult, this process can take days (Cockburn et al., 2022; Damen et al., 2021; Mesa et al., 2018). Delaminating cells have been likened to less-fit “losers” in a competition between basal cells; however, while loser cells delaminate during peak stratification, they die by apoptosis at earlier stages (Ellis et al., 2019). At a molecular level, the desmosomal adhesion protein Dsg1 has been linked to delamination *in vitro* by reducing tension at E-cadherin cell-cell junctions (Nekrasova et al., 2018). However, another study concluded that increased cell-cell adhesion via E-cadherin promotes delamination (Miroshnikova et al., 2018). These seemingly conflicting results could potentially be explained by the fact that the first study used immature (24h) raft cultures while the second used mature, fully stratified (6d) cultures, hinting at potential differences between early and peak stratification.

We conclude that delamination in the embryonic epidermis can be directed by HDs in a cell-autonomous manner, because all differentiative phenotypes are only observed in RFP+ cells. Our study shows that *Itgb4* is necessary for basal layer retention at E17.5, but dispensable at E15.5, a requirement that aligns with the developmental formation of HDs. Finally, our data indicate that the hyper-differentiative phenotype seen in Notch overexpression mutants can be partially explained by an increase in delamination propensity, through either direct or indirect control of integrin-β4 expression at the DEJ. Taken together, we provide the first evidence for the role of HD-laminin-332 adhesion in basal cell retention and differentiation.

## MATERIALS AND METHODS

### Animals

All mice in this study were housed in the AAALAC-accredited (000329), USDA registered (55-R-004), and NIH welfare-assured (D16-00256 (A3410-01)) animal facility at the University of North Carolina at Chapel Hill. All procedures and surgeries were conducted under IACUC-approved animal protocols (19-155, 22-121, 25-086). CD1 WT outbred mice (Charles River #022) were used for all fixed tissue imaging of lentiviral-infected embryos. For live explant imaging of the epidermis, mice were maintained on a mixed C57BL76/J background with alleles for *Krt14Cre; Rosa26B^mTmG^* (*Rosa26mTmG* (*Gt(ROSA)26Sortm4(ACTB-tdTomato,-EGFP)Luo/J*;Jackson Labs #007576) animals were used. All animals were infected with constructs described below using an *in utero* lentiviral injection. Note that all ages were defined as E0.5 the morning a vaginal plug was discovered. Additional developmental aging was performed by measuring the crown-rump length of embryos using a VEVO-2100 Ultrasound. Note that animals on the C57BL6/J background typically grew at a slower rate, roughly 0.5 days developmentally behind the CD1 strain mice.

### Lentiviral injections

The lentiviral injection procedures were performed according to Beronja et al. 2010., under the approved, aforementioned protocols (see “Animals”). Pregnant mice were anestized with 3% isoflurane for no more than 1hr and provided analgesics (5mg/kg meloxicam and 1-4mg/kg bupivacaine). One uterine horn was isolated, and 2-3 developing embryos were revealed at a time through a membrane, into a dish of sterile 1XPBS. Microinjections of 1.0-1.5uL lentivirus was delivered using an automatic pump into the amniotic cavity, visualized by ultrasound. Between 3-6 embryos were injected per pregnant dam, per surgery. Uninfected animals within the same litter were used as wild type, littermate controls per surgery. Following surgery, all embryos were returned into the body cavity of the dam, and surgical site was closed with sterile sutures and surgical staples. Mice were monitored for 4-7 days following surgery, and harvested at the appropriate developmental age, between E14.5 and E17.5.

### Constructs and RNAi

Short-hairpin RNAs (shRNAs) were generated targeting Integrin-β4, β1, and Laminin-α3 according to previously reported cloning protocols, described in Williams et al. 2011, 2014. NCBI Accession Numbers for a given shRNA clone were identified by gene name with the last four digits of the 21-nucleotide target sequence, for example *Itgb4^4124^*. shRNA constructs were cloned into vector backbones containing either the H2BRFP reporter sequence, or a puromycin selectable sequence. H2BRFP shRNAs were used for *in vivo* studies, while the puromycin selectable shRNAs were used to generate isogenic knockdown cell lines used for RT-qPCR. Both versions of the shRNA were packaged into lentivirus by using HEK 293FT cells, pMD2.G (Addgene #12259) and psPAX2 (Addgene #12260) helper plasmids. Due to reagent availability, some changes to the lentiviral production were made. First, plasmids were transfected with the Lipofectamine 3000 Transfection Kit, using Thermo-Fischer published protocols. To scale the lipofectamine reagents effectively, 358uL of Lipofectamine was added to 8.642mL of OptiMEM (for a total of 9mL). The 9mL of DNA mix and p3000 reagent were then added to two 500cm^2^ plates containing 81mL of OptiMEM media each. Secondly, Lonza UltraCulture Media was substituted for DMEM, 5% FBS, 2 µg/mL insulin, 1% L-glut, 1% sodium bicarbonate, 1% pen-strep, 1% sodium pyruvate in the viral production media (VPM).

### shRNA validation

Primary cultured basal keratinocytes were then infected with shRNA lentivirus on the puromycin containing vector. 48hrs following infection, cells were transferred from E-Low Media to E-Low Media with 2ug/mL puromycin and incubated for at least 7 days, with at least one passage during that time course. Selected cell lines were then lysed using the RNeasy Mini Kit (QIAGEN) to isolate RNA. cDNA was generated using the SuperScipt IV VILO Kit (11756050). RT-qPCR was then performed on dilute cDNA to determine mRNA knockdown efficiency using the TaqMan Fast Advance System on a ThermoFischer QuantStudio 6 Pro. The following TaqMan Probes were used: *Ppib* (Mm00478295_m1) and *Itgb4* (Mm01266840_m1). Cyclophilin B (*Ppib*) was used as the house keeping gene, and *Scramble* shRNA puro selected cell lines were used as the reference control. All material specifics and detailed protocols from the author are available upon request.

### Transmission Electron Microscopy (TEM)

To analyze epidermal ultrastructure, embryo heads from wild type littermate controls and infected animals were harvested and immediately placed into prewarmed TEM fixative (4% paraformaldehyde, 2.5% glutaraldehyde, 0.05M sodium cacodylate, pH 7.4, 5mM calcium chloride, and 1mM magnesium chloride) and incubated for 2hrs at room temperature. 2 mutants and 2 controls were submitted for processing for the early, E15.5 developmental time points, and 1 mutant and 1 control were submitted for the *Itgb4^4124^* infected epidermis E17.5 condition. Fresh fixative was used in each experiment, no more than 1 month old. Skulls were then left in 4C for no more than 3 weeks, until sample delivery to the UNC Microscopy Services Laboratory Electron Microscopy Team. Nape skin from the skulls was removed in small square pieces. The square nape skins were then resin embedded, and sagittal sectioned, and placed on TEM grids. Nape skin was collected as that is typically the most highly transduced area of tissue in the animal and therefore favors majority RFP+ cells. Only animals with high basal RFP+ infection by immunofluorescence were submitted for TEM analysis to aim for the most consistency in subsequent TEM analysis. During TEM imaging, we first identified the dermal-epidermal junction and captured reference images at 1.5kX and 8kX magnification for reference and to characterize gross architecture. Edges of the sample were not analyzed. ≥ 25 sequential images were taken at 30kX magnification at the dermal-epidermal junction. Representative images of hemidesmosomes and desmosomes from the lateral membranes were taken at 50-60kX magnification. Desmosomes across conditions appeared normal.

### Antibodies, immunohistochemistry, and fixed imaging

Integrin-β4 infected animals and their wild-type littermate controls were embedded whole into cassettes with Optimal Cutting Temperature (OCT, TissueTek) whole to avoid any damage to the presumed fragile epidermis. All tissue was then fresh frozen at -20C and stored at -80C for at least 24hrs prior to tissue sectioning. Infected animals were always mounted whole with a wild-type littermate to allow for direct comparison within the same slide for all immunohistochemistry. Frozen samples were then sectioned (10-12μm thick) on a Leica CM1950 cryostat. Staining of sectioned animals was conducted as previously described in Lough et al. 2019. Images were acquired using LAS AF software on a Leica TCS SPE-II, 4 laser confocal system on a DM5500 microscope. ACS Apochromat x40/1.5NA oil objectives were used for all fixed imaging.

The following antibodies were used Survivin (rabbit mAb 71G4B7, Cell Signaling 2808S, AB_2063948, 1:500), LGN (guinea pig, 1:500) (Williams et al., 2011), LGN (rabbit, Millipore ABT174, AB_2916327, 1:2000) (Williams et al., 2014), phospho-histone H3 (rat, Abcam AB10543, AB_2295065, 1:1000–5000), cleaved-caspase 3 (rabbit, Cell Signaling 9661, 1:1000), β4-integrin (rat, ThermoFisher 553745, AB_395027, 1:1,000), Cytokeratin-14 (chicken, Biolegend 906004, AB_2616962, 1:5000), Cytokeratin-10 (rabbit, BioLegend 905401, AB_2565049, 1:1000), GFP (chicken, Abcam AB13970, AB_300798, 1:1000), mCherry (rat, Life Technologies M11217, AB_2536611, 1:1000), RFP (rabbit, MBL PM005, AB_591279, 1:1000), Lamb3 aka Lam-5 (rabbit, Abcam, ab97765, 1:500). The following secondaries all raised in donkey, were diluted in gel block with normal donkey serum to reduce off target antibody staining: anti-rabbit AlexaFluor 488 (Life Technologies, 1:1000), anti-rabbit Rhodamine Red-X (Jackson Labs, 1:500), anti-rabbit Cy5 (Jackson Labs, 1:400), anti-rat AlexaFluor 488 (Life Technologies, 1:1000), anti-rat Rhodamine Red-X (Jackson Labs, 1:500), anti-rat Cy5 (Jackson Labs, 1:400), anti-guinea pig AlexaFluor 488 (Life Technologies, 1:1000), anti-guinea pig Rhodamine Red-X (Jackson Labs, 1:500), anti-guinea pig Cy5 (Jackson Labs, 1:400), anti-goat AlexaFluor 488 (Life Technologies, 1:1000), anti-goat Cy5 (Jackson Labs, 1:400), anti-mouse IgG AlexaFluor 488 (Life Technologies, 1:1000), anti-mouse IgG Cy5 (Jackson Labs, 1:400), anti-mouse IgM Cy3 (Jackson Labs, 1:500).

### EdU Injection and Analysis

To determine number of cycling basal cells, two independent dams were injected intraperitoneally with 5mM 5-Ethynyl-2-Deoxyuridine (EdU) in H2O following lentiviral surgery, 2hrs before harvest with 10uL per weight in grams. 3 mutant embryos and wild type litter mates were harvested from these two independent experiments. Secondary detection of EdU in tissue sections was performed by incubating tissue with ClickIt Chemistry (1XPBS, 200mM CuSO4 (final 2mM), 4mM Sulfo-Cyanine5 azide (final 8μM), 200mg/mL Ascorbate (final 20mg/mL) for 30 minutes following secondary antibody treatment. EdU-positive basal cells were then counted, and the percentage of EdU+, H2BRFP+ cells was determined relative to total H2BRFP+ basal cells, and likewise for RFP- and WT cells.

### Live Imaging

Live imaging was performed as described in previous publications, (see Patino Descovich et al., 2023, Lough et al. 2019, and Cetera et al., 2018). In short, epidermal explants from ages E14.5 to E16.5 were harvested from the midback of infected or uninfected *Krt14Cre; Rosa26B^mTmG^* animals. Explants were then sandwiched between a 1% Agar/F-Media patty (Ham’s F-12 Media, DMEM, 5% FBS, Sodium Bicarbonate, Sodium Pyruvate, and Pen/Step L-gluatmine) and a gas permeable membrane dish, (Lumox 3000). Confocal imaging was acquired using two microscopes, the Zeiss LSM-900 and the Andor Dragonfly spinning disk. The Andor Dragonfly microscope was funded with support from National Institutes of Health grant S10OD030223. On both microscopes, samples were maintained in a humidified, 37C temperature-controlled chamber, with 5% CO2. A 20X/0.80 Plan Apo and 40X/1.4 Oil Plan Apo objectives were used to image samples on the LSM-900. A Zyla Plus 4.2MP sCMOS camera and a HC PL APO 20x/0.75 LWD objective were used for imaging explants on the Dragonfly. For experiments on both microscopes, a 20-30µm z-stack was acquired every 5 minutes for between 2-6hrs. Imaging close to the epidermal edge was avoided due to edge disorganization and damage, limiting the effect wound healing may have on division orientation. Live imaging files were then deconvoluted using AutoQuantX 3.1. Individual image files of dividing cells were generated using the TimeSeriesAnalyzerv2.0, and division angles were measured in Fiji. Measurements of division orientation were taken at anaphase onset (t=0min) and anaphase completion (t=30mins). Imaging files are provided in the accompanied source data file. Additional imaging processing procedures are described in “Image Processing”.

### Data Analysis

#### Hemidesmosome Quantification

Hemidesmosomes (HDs) were identified in the 30kX magnification images. Density was quantified as number of HDs per (μm) length of tissue imaged. Sequential 30kX images had opposing, but not overlapping, views of the dermal-epidermal junction. Mature HDs were categorized based on being able to identify 1) electron density, 2) about 10nm in width, 3) had clear anchoring fibrils, and 4) had visible keratin filament extensions into the cytoplasm. Immature HDs were identified as being 1) electron dense but lacked in one or all the other identifiable features of mature HDs. Immature HDs also did cause noticeable depressions or flexing in the basal membrane of the basal keratinocytes, like mature HDs do. Density of all (Fig. 1H), only mature HDs (Fig. S1D) and only immature HDs (Fig. S1E) were determined as # HDs/μm per image (one dot) and plotted as a scatter plot. Additionally, the proportion of mature versus immature HDs was quantified as the fraction of total HDs identified. An Ordinary one-way ANOVA analysis was used to determine the statistically significant difference between WT and infected animal HD density in both mature and all HD analyses. A Chi-Square Analysis (Fisher’s exact test) was performed to determine the difference between mature and immature HDs between WT and infected animals.

#### Division orientation

Division orientation measurements were conducted on fixed and live imaging according to previously published methods, see Patino Descovich et al. 2023; Lough et al. 2019; Byrd et al., 2016; Williams et al., 2014; and Williams et al., 2011. In short, survivin was used as a midbody marker of dividing cells in telophase (late mitotic marker). The angle between pairs of daughter nuclei (labeled with DAPI, identified with survivin) was measured as a vector between the center of the DAPI signals and parallel to the basement membrane (Integrin-β4). In local areas of integrin-β4 depletion due to knockdown, the basement membrane angle was determined using residual integrin-β4 staining and the basal surface signal of Keratin-14. Cells in the hair placode were not calculated. Small z-stacks were taken from dividing cells to obtain the best imaging plane. All division angle vectors were calculated in Fiji, and ROIs are included in the raw data files. Live imaging division angles were measured in a similar fashion from individual frames over time, or at time points (t=0 and t=30). In RFP-cells, with no H2BRFP nuclear signal, nuclear positioning was estimated based on cell volume/membrane shape changes. Division angles ROIs and their corresponding measurements from live imaging files are included in the raw data files.

#### Basement Membrane Intensity

Basement membrane intensity of both integrin-β4 and laminin-β3 protein localization and intensity was quantified using a line profile analysis. *Itgb4* knockdown and control tissue were sectioned and stained on the same slides. They were stained for anti-integrin-β4 in AlexaFluor488, anti-mCherry Rhodamine Red X, and anti-Cytokeratin 14 in Cy5. On the same day, using the same laser power settings, gain and offset, 6 or more images were taken from anterior to posterior of the mouse. In Fiji, 10-pixel thick lines were drawn according to the shape of the basal surface of Cytokeratin-14 staining corresponding to either RFP+ zones, RFP-zones (at least 3 cells in a row), and in zones from WT animals. The sum fluorescence intensity for each point along the line (averaged from the 10-pixel width) was then divided by the total length of the line measured. The intensity per RFP+, RFP-, and WT zone were normalized to the average of the WT zone for that corresponding wild type littermate, such that the average of WT intensity was 1 (100%). As laminin-β3 is secreted, and likely would not show regional differences in intensity like integrin-β4, the entire length of the dermal epidermal junction in control and *Lama3^1436^*was measured.

#### LGN & Pericentrin Localization

Localization of LGN crescents (eg. Apical, non-apical (ie. basal, lateral), unpolarized, absent) were categorically determined from PHH3 positive basal cells. Chi-square analyses were used to determine the difference in count across the categories for RFP+, RFP-, and WT basal cells. Pericentrin localization was determined by measuring the angle between the pericentrin puncta, bisecting the nucleus, relative to/parallel to the basement membrane. In areas of integrin-β4 local depletion, Keratin-14 or laminin-β3 signal was used to determine the basal surface of the tissue. Angles from individual cells across multiple animals was represented as a scatter plot with angles ranging from 0-90^0^ from the basement membrane.

#### Spinous Layer Thickness

Spinous layer thickness was determined from mutant and control tissue sectioned and stained on the same slides. Tissue was stained for anti-Cytokeratin 10 in AlexaFluor488, anti-mCherry and anti-Integrin β4 in Rhodamine Red X, and anti-Cytokeratin 14 in Cy5. All the animals were sampled with 1µm optical sections for a total volume of 15μm on the 40Xoil objective of the SPE 5500D microscope. Each epidermis was imaged along the anterior-posterior axis in at least 5 positions. Infected animals were only imaged in areas of relatively high basal cell infection. Images were then processed in Fiji using a Macro with the following operations: 1) generate maximum projection of only the Cytokeratin 10 channel, 2) threshold using Huang metrics, 3) “Fill holes” of the mask, 4) use “open” function to eliminate background pixel contribution, 4) select and measure mask area. All spinous thickness mask TIFFs are saved and included in the raw data included. Mask TIFFs were then traced along the basal surface to determine the length of the epidermis measured. Total area of the spinous layer was then normalized to the length of the epidermis measured. Data is represented as the average of the multiple positions along the A-P axis per animal.

#### Delamination Quantification: K10, K14 Dual Positivity

To determine rate of delamination from fixed samples, tissue stained in the same way as the spinous layer thickness quantification was imaged at least 6 times from anterior to posterior. Using the Fiji plugin “Cell Counter” 6 categories of cells were determined: 1) Total basal cells (K14 and DAPI), 2) Dual positive (K14, K10 and DAPI), 3) Total infected basal cells (K14, H2BRFP, and DAPI), 4) Dual positive cells with clear basement membrane contact (K14, K10, DAPI, full or reduced contact (ie. upside-down, tear-drop shaped), 5) Total spinous layer cells (K10, DAPI), and 6) infected spinous layer cells (K10, H2BRFP, and DAPI). Then totals for each category were determined and represented as a fraction over that total population: RFP-delamination % = RFP-Dual Positive with Contact/Total RFP-basal cells. RFP+ delamination % = RFP+ Dual Positive with Contact/Total RFP+ basal cells. Basal cell infection % =Total RFP+ basal cells/Total basal cells. Each dot represents the percent dual positive basal cells per condition for an animal, where at least 45 basal cells per condition were counted for each animal. This threshold was intended to reduce the effect size of dual positive events over a small population. All cell count files are saved and included in the raw data file.

#### Delamination Quantification: Differentiation Index

Based on the cell count categories from the delamination index quantification, we can determine the percentage of spinous layer cells that are H2BRFP positive relative to the basal layer cells that are H2BRFP positive. Occupancy of the H2BRFP signal is a pseudo-lineage trace system because all cells exposed to lentiviral shRNA are basal at the time of infection. We then determined the proportion of DAPI cells that were RFP+ in the respective layers, where % spinous RFP+ was divided by % basal RFP+ to give a differentiation index score generally between 2.5 and 0.5. A differentiation index of 1.0 corresponds with equal infection rates in each layer, indicating no occupancy preference.

### Statistical analyses and graphs

All measurements were recorded into Microsoft Excel Worksheets per experiment or genetic background uploaded to GraphPad Prism v10, where graphs and statistical analyses were performed. Error bars represent standard error of the mean. Statistical tests of significance were determined by the following tests, Student’s T-test for two averages, One-Way ANOVA for comparison of 3 or more averages, Fischer’s exact test/Chi-Square for comparing distributions of categorical data, and observed versus expected/Chi-Square analysis for comparing telophase correction outcomes in Fig. 3D-F. Figures were generated from Prism graphs uploaded to either Adobe Illustrator as EMF files. Sample sizes and cell numbers were chosen based on the power analyses conducted for similar studies in Patino Descovich et al., 2023; Lough et al., 2019; and Williams et al. 2014. All raw data is available for closer examination in the included data file.

## Acknowledgements

The authors would like to acknowledge Williams lab members: Amber Altrieth, Lauren Griffith, Carlos Patino Descovich, and Bethany Brown for their insightful feedback, technical support, and manuscript editing. Joint laboratory meeting discussion between the Peifer, Bergstrahl-Finegan, and Lovegrove lab have also been instrumental in the organization of the manuscript. We are immensely grateful to Pablo Ariel, Kristen White, and Jillann Madren in the Microscopy Services Laboratory (MSL) core facility for light microscopy assistance and TEM tissue preparation and imaging. JK also thanks AMV and ABKV for support and transportation during experimentation and manuscript preparation.

## Funding

National Institutes of Health, R01 AR077591, Scott E Williams

United States – Israel Binational Science Foundation, 2019230, Scott E Williams

National Institutes of Health, F31 DE033915-01, Juliet S. King

University of North Carolina at Chapel Hill Cell Biology and Physiology Department, T32 GM133364, Juliet S. King

