## Supplemental Data for "Hemidesmosomes regulate epidermal differentiation during embryogenesis"

**SUPPLEMENTAL FIGURES & SUPP. FIGURE LEGENDS**

**Supp. Figure 1. Mosaic loss of Itgb4 alters embryonic basement membrane organization without overt blistering** (A) Relative mRNA abundance of *Itgb4* transcripts from *Scramble*, $Itgb4^{4124}$, and $Itgb4^{2326}$ infected, puromycin-selected primary keratinocytes. mRNA levels were normalized to *Scramble* levels per technical replicate, represented by individual dots. (B) Representative images of control and $Itgb4^{2326}$ infected epidermis stained with integrin-β4 (green), H2BRFP (magenta), and DAPI (blue). Note: blue arrowheads indicate uninfected regions. (C) Intensity of the integrin-β4 staining in WT (black), RFP- (grey), and RFP+ (light blue) regions in $Itgb^{2326}$ mutant epidermis. Intensity is normalized to the average of the WT integrin-β4 intensity. Individual dots represent measurements across 3 control and 3 infected animals from two litters. (F) Mature only HD density in control and $Itgb4^{4124}$ mutants. (G) Immature only HD density in control and $Itgb4^{4124}$ mutants. Note that each dot represents a sequential position along a TEM micrograph for 1 control and 2 mutants at E15.5, and 1 control and 1 mutant at E17.5. Scale bars indicate 25μm in (B). p-values: ns, not significant; * $p<0.05$, ** $p<0.001$, **** $p<0.0001$.

**Supp. Figure 2. *Itgb4* loss leads to decreased basal occupancy but does not impact division orientation or polarity.** (A-B) Representative image of phospho-histone H3 positive (green) basal cell (DAPI) in an $Itgb4^{4124}$ infected epidermis (H2BRFP, magenta), with quantification (*B*) of number of PHH3+ basal cells per total basal cell population. (C-D) Representative image of EdU+ (green) basal cell (DAPI) in wild type littermate and an $Itgb4^{4124}$ infected littermate epidermis (H2BRFP, magenta), with (*D*) quantification of number of EdU+ basal cells per total basal cell population, where each dot is the percentage of EdU+ basal cells for that population per animal, across 2 independent litters. (E-F) Representative image of cleaved caspase-3 positive (green) basal cell (DAPI, blue) in a mutant $Itgb4^{4124}$ mutant epidermis (H2BRFP, magenta) with (*F*) quantification of the total number of cleaved caspase-3 positive basal cells observed across the entire back skin sagittal section. Three epidermal back skin samples were analyzed for 3 $Itgb4^{4124}$ mutant animals, harvested from three independent LUGGIGE litters. Individual dots correspond to the average number of cleaved caspase-3+ positive basal cells observed across the 3 sagittal sections. The number of cleaved caspase-3+, RFP+ cells is reported as a fraction next to the average for that mutant animal. (G) Cumulative frequency plots of division orientation for control (black) and *Itgb4*$^{4124}$ (orange) infected animals at E16.5-17.5. Same graph as in Fig. 2F, but includes the RFP- cumulative frequency distribution line (grey). (H) Cumulative frequency plot of division orientation for second hairpin, *Itgb4*$^{2326}$ RFP+ cells (light blue), WT (black), and RFP- (grey). (I-J) Par3 (green) localization in wild type littermate and *Itgb4*$^{4124}$ mutant (H2BRFP, magenta) epidermis, with basal layer marked in K14 (white). (K) Relative enrichment of Par3 across the cortex of the cell, where 50% of the normalized distance corresponds to the apical-most region of the cell. Line profiles were traced from the bottom of the left lateral membrane to the bottom of the right lateral membrane for wild type and infected basal keratinocytes. p-values: ns, not significant.

**Supp. Figure 4. Loss of integrin-β4 or its ligand laminin-332 cell-autonomously increases delamination.** (A) Scatter plot of percentage of dual positive basal cells per WT (black), RFP+ (magenta) and RFP- (grey) population in *Itgb4*$^{4124}$ mutant epidermis at pre-stratification timepoint, E15.5. Each dot is the total percentage dual positive per animal. (B) Representative images of wild type and *Itgb4*$^{4124}$ littermates stained with Laminin-β3 (green, black in single channel), H2BRFP (magenta), and DAPI (blue). (C) Normalized spinous layer thickness in $\mu m^2$ in wild type and *Lama3*$^{1436}$ mutant littermate epidermis at E16.5 and E17.5. (D) Whole mount immunofluorescent images of *Lama3*$^{1436}$ epidermis on a *Krt14*$^{Cre}$*; Rosa26B*$^{mTmG}$ background with the basal *en face* view (single z-slice) on the left and the spinous and periderm layers (single z-slice) in view on the right. Note the rare H2BRFP positive cells in the basal layer with increased number of H2BRFP+ nuclei in the spinous and periderm layer. p-values: ns, not significant.

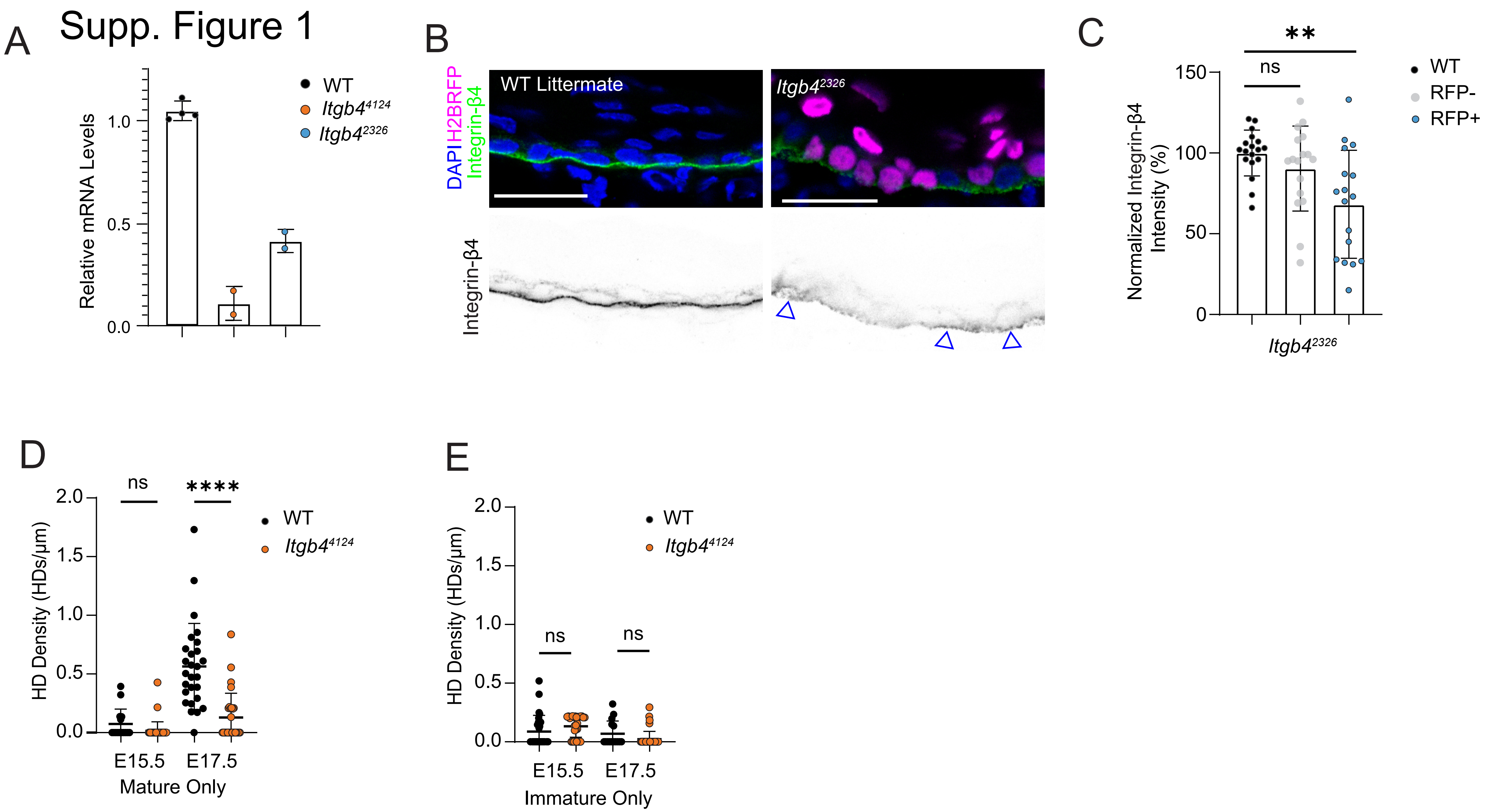
Supp. Figure 1
A
Relative mRNA Levels
1.0
0.5
0.0
WT
Itgb4^4124
Itgb4^2326
B
DAPI H2BRFP Integrin-β4
WT Littermate
Itgb4^2326
Integrin-β4
C
Normalized Integrin-β4 Intensity (%)
150
100
50
0
ns
**
WT
RFP-
RFP+
Itgb4^2326
D
HD Density (HDs/µm)
2.0
1.5
1.0
0.5
0.0
ns
****
WT
Itgb4^4124
E15.5
E17.5
Mature Only
E
HD Density (HDs/µm)
2.0
1.5
1.0
0.5
0.0
ns
ns
WT
Itgb4^4124
E15.5
E17.5
Immature Only

### Supp. Figure 2

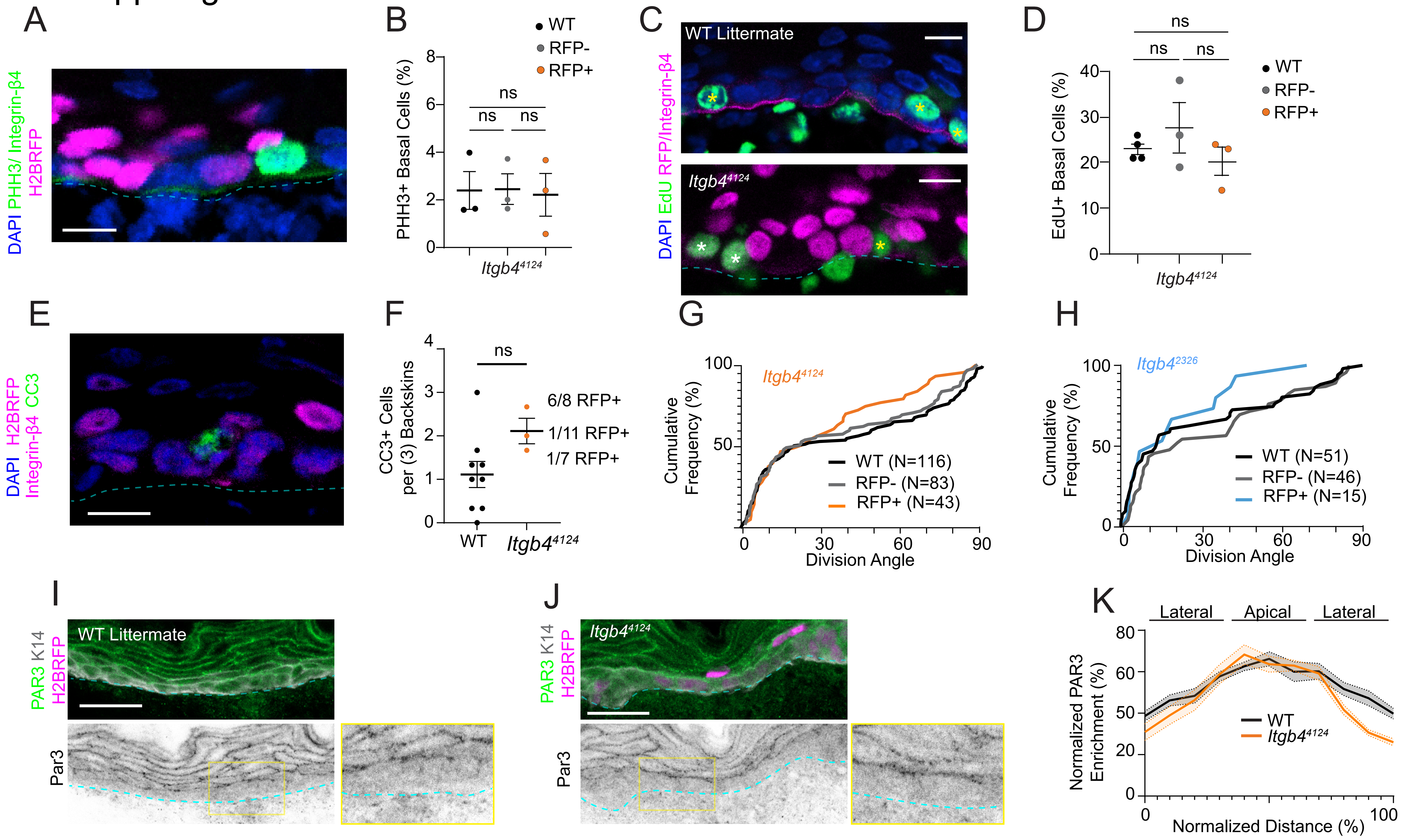

### Supp. Figure 4

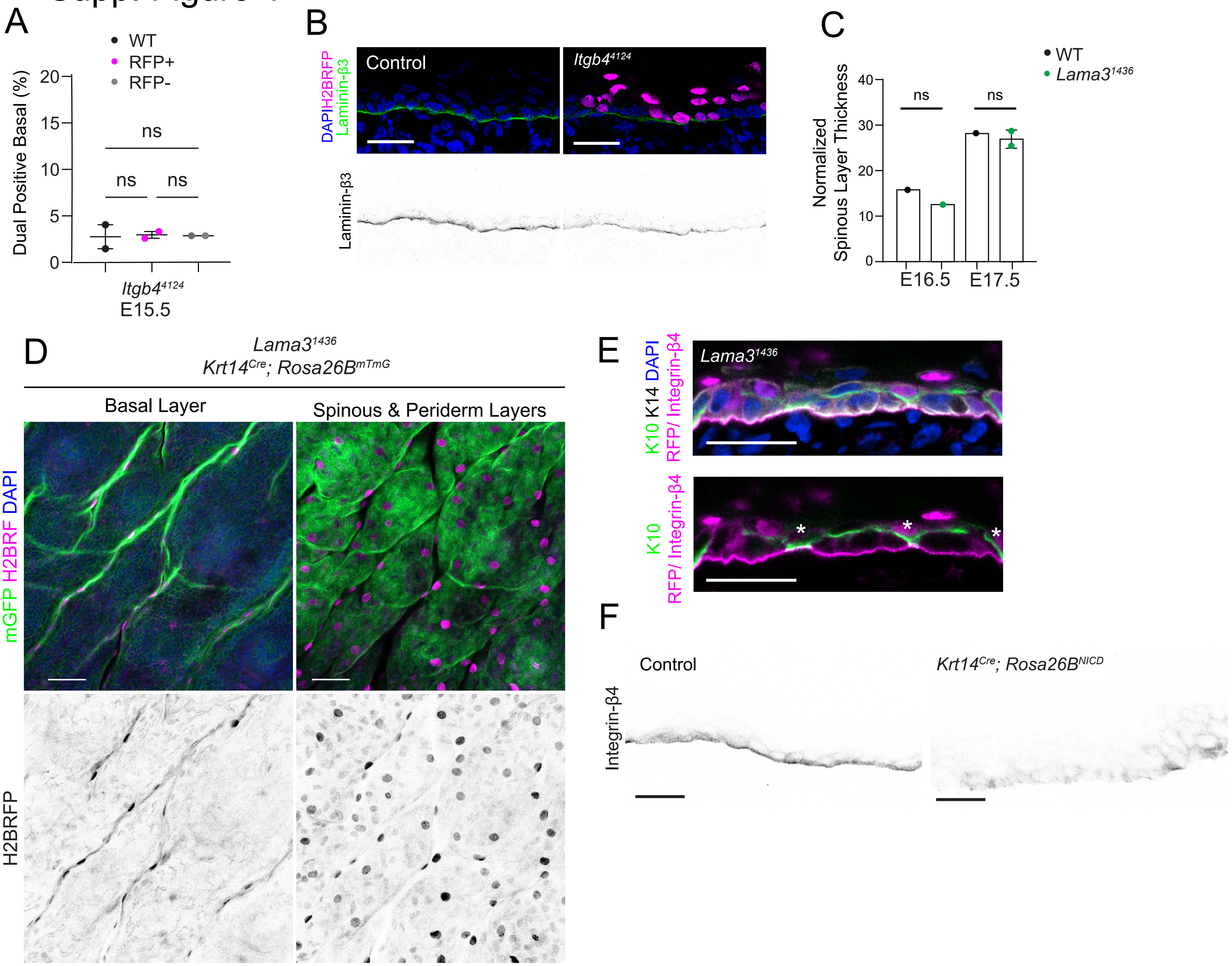
